# Quantifying visual acuity in *Heliconius* butterflies

**DOI:** 10.1101/2023.07.10.548283

**Authors:** Daniel Shane Wright, Anupama Nayak Manel, Michelle Guachamin-Rosero, Pamela Chamba-Vaca, Caroline Nicole Baquet, Richard M. Merrill

## Abstract

*Heliconius* butterflies are well-known for their colourful wing patterns, which advertise distastefulness to potential predators and are used during mate choice. However, the relative importance of different aspects of these signals will depend on the visual abilities of *Heliconius* and their predators. Previous studies have investigated colour sensitivity and neural anatomy, but visual acuity (the ability to perceive detail) has not been studied in these butterflies. Here, we provide the first estimate of visual acuity in *Heliconius*: from a behavioural optomotor assay, we found that mean visual acuity = 0.49 cycles-per-degree (cpd), with higher acuity in males than females. We also estimated visual acuity from eye morphology and reported slightly lower values (mean visual acuity = 0.38 cpd), but acuity was still higher in males. Finally, we estimated how visual acuity affects *Heliconius* visual perception compared to a potential avian predator. Whereas the bird predator maintained high resolving power, *Heliconius* lost the ability to resolve detail at greater distances, though colours may remain salient. These results will inform future studies of *Heliconius* wing pattern evolution, as well as other aspects in these highly visual butterflies, which have emerged as an important system in studies of adaptation and speciation.

## 1. Introduction

Since Bates (1) first described mimicry over 160 years ago, studies of *Heliconius* butterflies have made an important contribution to our understanding of adaptation and speciation (2). These Neotropical butterflies are well known for their diversity of bright colour patterns, which both advertise distastefulness to potential predators (3–8) and are used during mate choice (9). Because males distinguish between the warning colour patterns of con- and hetero-specific females (and to some extent against con-specific females from populations with different patterns), colour pattern contributes an important premating reproductive barrier, e.g. (10–12). *Heliconius* also use visual cues during foraging and host plant selection (13,14). Vision, therefore, plays a crucial role in *Heliconius* behaviour, and studies of *Heliconius* have increasingly considered vision, especially with respect to colour perception (15–19) and neuroanatomy (20,21). However, a key element of *Heliconius* visual ecology has not yet been studied, specifically visual acuity.

Visual acuity is the ability to perceive detail in a visual scene and is one of the three fundamental parameters of visual systems (the others being spectral sensitivity and temporal resolution (22,23)). Visual acuity is typically reported as the number of black and white stripe pairs that an organism can discriminate within a single degree of visual angle (cycles-per-degree; cpd) (23). Across animal taxa, acuity varies greatly. For example, human visual acuity is 72 cpd (24), whereas fruit fly visual acuity is only 0.09 cpd (25). This discrepancy is particularly relevant for researchers designing studies of visual signals, whereby hypotheses about the form and function of a given trait may not consider the visual capacity of the receiver (26). Caves, Frank and Johnsen (27) demonstrated this principle empirically by showing that cleaner shrimp visual acuity is too poor to resolve colour patterns previously believed to be used in intraspecific signalling. This result nicely illustrates the potential downfalls of biassing studies towards our own perceptual abilities.

A battery of studies have left little doubt that *Heliconius* males can distinguish between potential mates with divergent colour patterns (6,10–12,28–30). However, visual acuity has never been measured in these butterflies, so it is unclear how well the butterflies can perceive these signals at different distances, or what is the relative importance of colour vs. pattern at different points of courtship (e.g. long-range vs short-range attraction). For other diurnal butterflies, visual acuity has been reported from 0.66 cpd (*Colias eurytheme* (31)) to ∼1.0 cpd (*Morpho peleides* (32)), and as body size scales with eye size, visual acuity tends to increase in larger butterflies (33). *M. peleides* is larger than *Heliconius*, so visual acuity values <1.0 cpd are expected for *Heliconius*. If this is true, the ability of *Heliconius* to resolve wing pattern details is questionable, particularly at greater distances. Avian predators, on the other hand, likely perceive the same colour patterns clearly - *Heliconius* bird predators are unknown, but visual acuity studies in other insectivorous birds suggest high visual acuity (e.g. *Acanthiza chrysorrhoa* = 25.6 cpd (34); *Zosterops lateralis* = 18.5 cpd (35)).

Here, we measure visual acuity in *Heliconius* butterflies for the first time. We took two complementary approaches. First, we used a behavioural assay to measure optomotor responses, where animals turn in the direction of a rotating stimulus so as to minimise displacement of the moving image (25). Second, we used an anatomical approach to quantify the number of photoreceptors, allowing estimates of visual acuity from eye morphology. By more thoroughly investigating the potential mismatch between the visual abilities of *Heliconius* and their predators, our results shed light on the evolution and function of the warning patterns.

## 2. Methods

### (a) Study species

We established a stock of *Heliconius erato cyrbia* from wild individuals caught in forests near Balsas (3°43′60′′S, 79°50′45′′W) in Southern Ecuador. These were maintained at the Universidad Regional Amazónica IKIAM in Tena, Ecuador, and replenished with wild individuals intermittently over the course of the experiment. All butterflies were maintained under common garden conditions in outdoor insectaries in 2 x 2 x 2.3 m cages, where they were provided with 20% sugar solution, and *Lantana* sp. and *Psiguria sp*. flowers as a source of pollen. Eggs were collected regularly from the hostplants, *Passiflora punctata*, provided in the insectaries. The larvae were reared individually in pots and fed with fresh leaves from the host plants. All butterflies were marked with unique identification codes on their wings after eclosion.

### (b) Behavioural estimates of visual acuity

An optomotor device (Fig. S1) was built following Caves et al. (36), which permits a non-invasive and reliable behavioural assay for studying visual acuity across taxa. Briefly, the device consisted of interchangeable visual stimuli of alternating vertical black and white stripes printed on waterproof paper (145 μm, Premium NeverTear, Xerox, CT, USA) on a rotating wheel around a fixed white PVC base (17 cm radius). The width of one cycle (a set of alternating black and white stripes) was calculated as *cycle width (mm) = [(C/360) / a]*, where ‘*C*’ is the circumference of the experimental arena and ‘*a*’ is the intended visual acuity in cycles-per-degree (cpd) (36). We used stimuli with spatial frequencies of 0.3 cpd to 1 cpd (in pilot trials, butterflies consistently responded to cpd levels >0.3). As visual acuity depends on the distance between the stimulus and the perceiver, all butterflies were restrained in a clear PLEXIGLAS cylinder (4 cm radius; 15 cm height) at the centre of the base. A blank stimulus was used to confirm that responsiveness was due the moving stripes and no other external cues.

All assays were conducted inside (at room temperature) at IKIAM University, illuminated by an overhead LED ring lamp, and video recorded from above. Butterflies (10+ days post eclosion) were tested only when they stopped crawling on the cylinder, followed by 3-4 rotations of the stimulus (alternating between clockwise and counter-clockwise for 10 seconds each, at 3 rpm). A positive response was scored only if the butterfly changed the orientation of its head/antenna in the direction of the moving stimulus in two consecutive stimuli rotation reversals (see supplemental video S1). All positive responses were confirmed from videos, and butterflies with unclear responses were excluded from the experiment.

Male butterflies tend to have larger eyes than females (33), so we explored how visual acuity differs between male vs. female *H. e. cyrbia*, while accounting for the time-of-day butterflies were tested (10:00∼16:00) and potential observer and outside weather effects (all trials were conducted indoors, but the butterflies were kept in outdoor insectaries) using generalised linear mixed models implemented in the lme4 package in R (37). The generalised linear model (family = Gamma) was as follows: *glmer(visual acuity) ∼ sex + time + (1*|*observer) + (1*|*weather)*. The significance of fixed effect parameters (*sex* and *time*) was determined by likelihood ratio tests (LRT) via the *drop1* function and minimum adequate models (MAM) were selected using statistical significance (38,39). Model assumptions were confirmed via visual inspection (residual vs. fitted and normal Q-Q plots). We used the *Anova* function in the car package (40) to estimate the parameters of significant fixed effects based on LRT.

### (c) Morphological estimates of visual acuity

The insect compound eye consists of numerous independent photosensitive units, ommatidia, each of which receives visual information and transfers it to the brain. Variation in ommatidial number/density directly affects visual acuity (25,41,42), which can be estimated from ommatidia counts (see below).

Specimens from the stock population were preserved in DMSO/EDTA/NaCl (43) and stored at -20° C. Following previously published methods (44), frozen specimens were thawed at room temperature and both eyes were removed and placed in 20% sodium hydroxide (NaOH) for 18-24 hours to loosen the tissues behind the cuticular cornea. The following day, the cuticle was cleaned of excess tissue, mounted on a microscope slide in Euparal (Carl Roth GmbH, Germany), and left to dry overnight.

We used ImageJ/Fiji (45) to analyse each mounted cornea for the total number of ommatidia. All slides were imaged at 7.5x on a Leica M80 stereomicroscope fitted with a Leica Flexacam C1 camera and the Leica Application Suite X (LAS X) software. Each image contained a 1 mm scale bar to calibrate the measurements. Ommatidia counts were measured via image thresholding and the *Analyze particles* function (methods provided as supplementary information). To account for differences in body size, the hind legs of each butterfly were also removed and imaged; hind tibia length was measured using the *Straight line* and *Measure* options.

In a compound eye hemisphere covering 180° of space, the number of ommatidia ‘*n*’ that can cover the hemisphere without overlap is *23818/(ΔΦ*^*2*^), where *ΔΦ* is the interommatidial angle (25). Thus, the number of ommatidia (*n*) can be used to calculate the interommatidial angle as: *ΔΦ = (23818/n)*^*1/2*^, and from this, visual acuity can be estimated as: *acuity = 1 / (2ΔΦ)* (equation 1 from (25)). Based on the strong correlation between the left and right eye ommatidial counts (r = 0.949, t = 8.01, df = 7, p < 0.001), only one eye from each individual was used for calculating visual acuity. For uniformity, we always used the left eye unless it was damaged or imaged poorly, in which case the right eye was substituted.

We used a generalized linear model (glm) to explore how the morphological estimates of visual acuity differs between the sexes while accounting for body size (hind tibia length) as g*lm(visual acuity ∼ sex + tibia length)*. We also investigated sex-specific differences in eye morphology as l*m(log10(ommatidia count ∼ sex + log10(tibia length))*. We used log10-transformations to normalise the residuals around the allometric relationship between ommatidia count and tibia length (46). Model simplification and parameter estimates were as detailed above.

### (d) Bird-butterfly comparison

We used the AcuityView package in R (26) to estimate how the visual acuity values reported here influence *H. e. cyrbia* perception. We also modelled how bird predators may view the same visual scene. The exact bird species that prey upon *Heliconius* are unknown, as are their visual acuity values, but prior studies (e.g. (47)) have used insectivorous Passerines as representative predators. Thus, we used the mean acuity value (22.05 cpd) of two insectivorous Passerines (*Acanthiza chrysorrhoa and Zosterops lateralis* (34,35)) as a proxy.

## 3. Results

### (a) Behavioural estimates of visual acuity

We tested 26 males and 23 females in the optomotor assay. Overall, the mean behavioural visual acuity (± standard error) for *H. e. cyrbia* was 0.491 ± 0.027 cpd. Mean male visual acuity was 0.547 ± 0.043 cpd, which was significantly higher than the mean value for females (0.427 ± 0.023; χ^2^ = 5.35, df = 1, p = 0.021; Fig. 1A, Table 1). The time of day the butterflies were tested did not influence the results (p > 0.4)

**Table 1.** Visual acuity estimates (mean cycles-per-degree ± standard error) for *H. e. cyrbia* males and females from the behavioural optomotor assay and morphological analyses.

|  | Behavioural | Morphological |
| --- | --- | --- |
| **male** | $0.547 \pm 0.043$ | $0.384 \pm 0.005$ |
| **female** | $0.427 \pm 0.023$ | $0.373 \pm 0.003$ |

**Figure 1.**
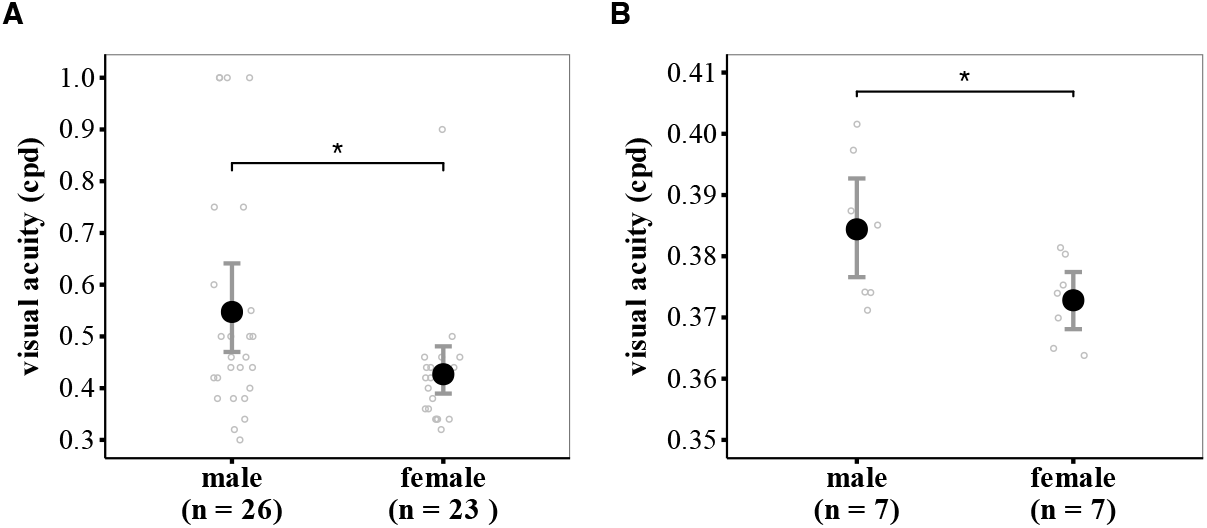
Estimates of visual acuity in the **(A)** behavioural optomotor assay and from **(B)** morphological analyses both indicate that male *H. e. cyrbia* have higher visual acuity (in cycles-per-degree, cpd) than females. Solid black circles represent the mean values, and grey error bars show 95% confidence intervals. * indicates p < 0.05.

### (b) Morphological estimates of visual acuity

We analysed the eye morphology of 7 males and 7 females. The morphological estimates of visual acuity were lower than the behavioural results presented above (see Table 1); the overall mean was 0.379 ± 0.003 cpd, compared to 0.491 cpd in the behavioural assay. Nonetheless, patterns were similar, and males had significantly higher visual acuity than females (χ^2^ = 4.92, df = 1, p = 0.027, Fig. 1B, Table 1). This can be attributed to the fact that males had more ommatidia than females (F_1,12_ = 4.93, p = 0.046), despite no significant differences in body size (using hind tibia length as a proxy; p > 0.7).

### (c) Bird-butterfly comparison

Using the mean visual acuity values from the optomotor assay, we found that *H. e. cyrbia* has relatively poor visual resolution, particularly at larger distances (Fig. 2). In contrast, our proxy bird predator maintained high resolution at all distances.

**Figure 2.**
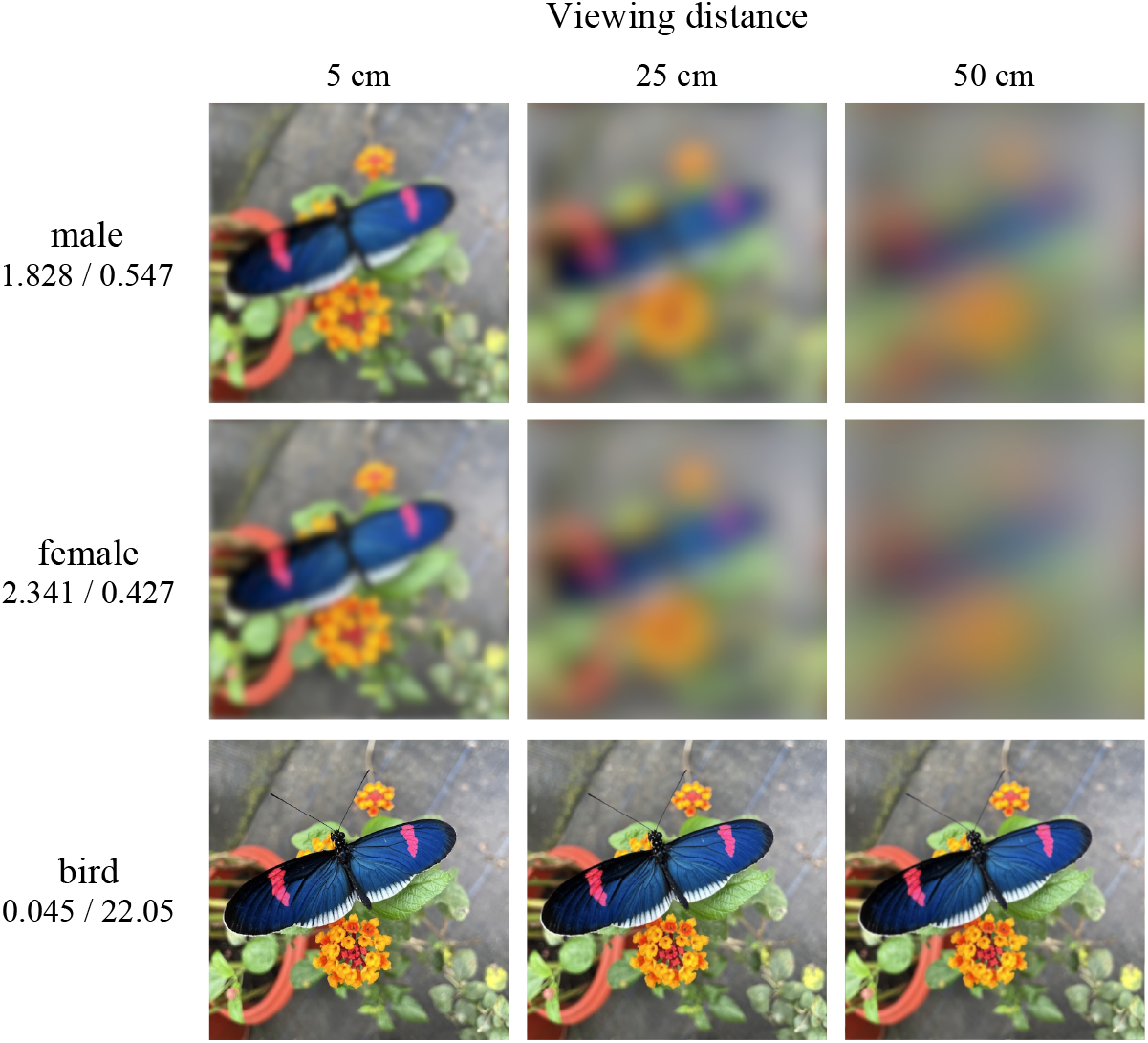
Perceptual estimates of *H. e. cyrbia* males (top) and females (middle) using the mean visual acuity values from the optomotor assay (generated with the AcuityView R package (26)). For comparison, the bottom row presents the perceptual estimate of a hypothetical bird predator viewing the same visual scene. *Heliconius* bird predators and their visual acuity values are unknown, so we used the mean visual acuity value of two insectivorous Passerines (*Acanthiza chrysorrhoa and Zosterops lateralis*; (34,35)). Values on the left side represent the minimum resolvable angle in degrees / visual acuity in cycles-per-degree.

## 4. Discussion

There are three fundamental parameters of visual systems: spectral sensitivity, temporal resolution, and visual acuity (22,26). For *Heliconius* butterflies, spectral sensitivity is well characterised (15–19), but temporal resolution and visual acuity are undocumented. Here, we report the first measures of visual acuity in *Heliconius* using two approaches: mean behavioural visual acuity was 0.49 cpd, whereas morphological estimates were somewhat lower (0.38 cpd). Our use of two experimental approaches provides insight into the morphological correlates of visual acuity in *Heliconius*, while also accounting for post-retinal processing involved in visual perception and behavioural output. We report lower morphological estimates of visual acuity than in the behavioural assay, but the morphological estimates do not account for higher-order visual system processing. Despite this discrepancy, both approaches revealed higher visual acuity in males than females, as has been observed in other butterflies (33). Morphological and ecological differences between the sexes might explain this sexual dimorphism; males have more ommatidia (larger eyes) than females, despite similar body size (see results above), and males actively search for and identify conspecific mates (48,49).

*Heliconius* wing patterns warn potential predators that these butterflies are unprofitable prey and are also used during mate choice. Previous studies have repeatedly demonstrated that *Heliconius* males can distinguish between potential mates with divergent colour patterns both between (10,11,29), and to a lesser extent, within species (6,12,28,30). However, our results suggest that these signals may not be viewed equally by conspecifics and predators. Although visual acuity will always limit what information can and cannot be perceived (26), our visual representations (Fig. 2) are of course only approximations informed by acuity measurements and cannot depict what the animals actually see. Other factors, such as neural image enhancement and motion detection are involved, but these processes cannot add information to the visual image (26). Nevertheless, our recreation of how the butterflies and a potential avian predator may view the same visual scene highlight that while potential predators maintain high visual resolving power at all tested distances, *Heliconius* will quickly lose the ability to resolve pattern detail as distances increase.

In contrast to specific fine-scale patterns, colour elements may remain salient for the butterflies across a broader range of distances, suggesting that colour may be more useful for e.g. long-range attraction. This is consistent with experimental work showing that *Heliconius* colours likely have a greater influence than patterning for both predators and conspecifics (6), and that more prominent shifts in male preference are often associated with major colour differences (e.g. white to yellow, or white to red shifts in forewing colouration (50)). Such differences between avian and *Heliconius* perceptive abilities may help to inform future studies of *Heliconius* wing pattern evolution. While we can never fully appreciate the perceptive abilities of our study animals, our results highlight differences that our own perceptual biases may have otherwise missed in an important model of adaptation and speciation, which relies heavily on visual information to survive and reproduce.

## Acknowledgements

We are grateful to the Ministerio del Ambiente, Agua y Transición Ecológica for permission to collect butterflies in Ecuador (MAATE-DBI-CM-2021-0176; MAATE-DNB-CM-2021-0176). Lucie Queste, Gladis Grefa, José Borrero Malo, Sophie Smith, Whitney Wright, Tamia Chimba, Esteban Arévalo-Morocho and Samantha Vasco Viteri contributed to plant and butterfly stock maintenance. Lisa Ammer assisted with image analyses. This research was supported by an ERC starting grant (Grant number: 851040) to RMM.

