## Supplemental information for "Quantifying visual acuity in *Heliconius* butterflies"

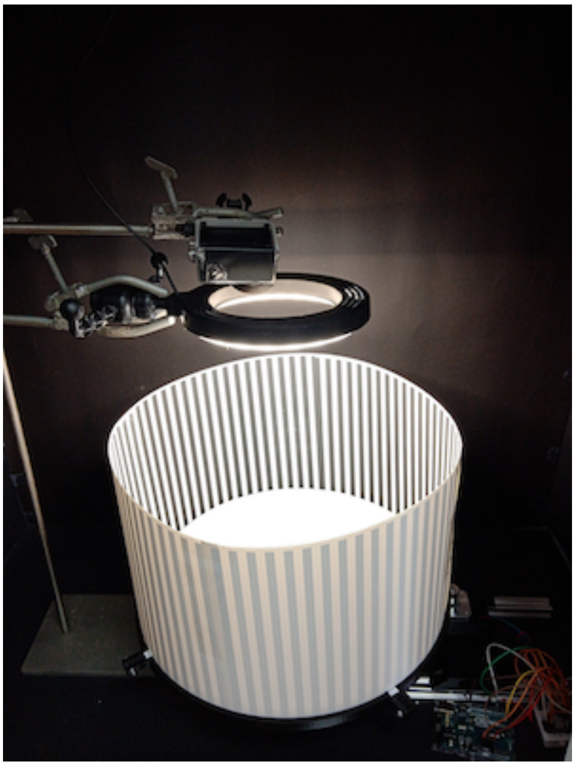

**Figure S1.** The optomotor set up. Butterflies were contained at the centre of the fixed white PVC base in a clear PLEXIGLAS cylinder (not pictured) while the visual stimulus (alternating black and white lines) rotated (in both directions for 10 seconds at 3 rpm). All trials were illuminated with a LED ring lamp and video recorded from above (see also: video S1).

### Image analysis methods

#### Facet count:

1) Open ImageJ/Fiji.

2) Open the desired image via *File* → *Open* or by dragging the image onto the console.

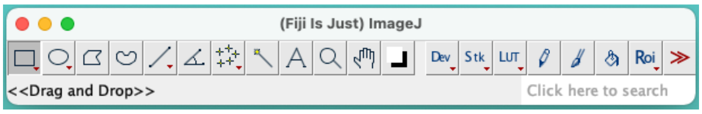


3) *Analyze* → *Set Scale* to specify the **distance in pixels** per **unit of length**. This is can be assessed via the line tool in the console.

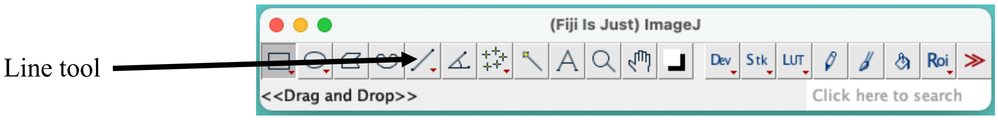


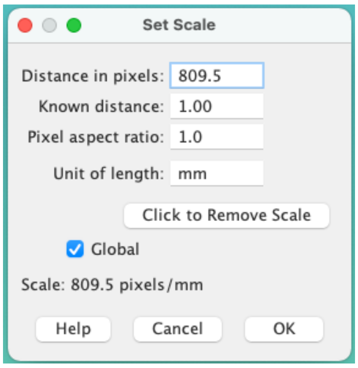


   a. Click and drag a line the length of a known distance in the image, e.g., the scale bar, and then measure with *Analyze* → *Measure*.
   b. Enter this value as the **Distance in pixels**, specify the **Known distance** (of the scale bar), and the **Unit of length**.
   c. Select **Global** to apply this to all pictures opened in the current session.
      <u>Note</u>: these values will need to be reassessed for images taken at different zoom/focus levels.

4) *Image* → *Type* → *16-bit* (8-bit is also ok). The image should change to grey-scale.

5) *Image* → *Adjust* → *Threshold*.
   a. Use the sliders to adjust the threshold so that the entire eye/cuticle is highlighted (i.e., move the bottom slider to the right to increase the threshold, if necessary).

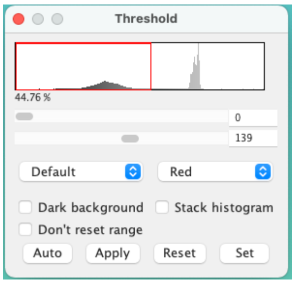


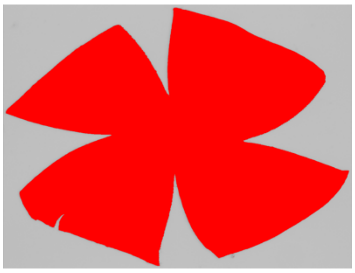

6) *Process* → *Subtract background.*
   a. Rolling ball radius: 50 pixels
   b. No additional options selected
   c. Click *OK* (the image will darken)
   d.

7) Use the threshold sliders (the window should still be open) to adjust the threshold so that only the individual facets are highlighted.

a. This is best achieved by first moving the top slider to the right and then the bottom slider to the left.

b. The goal is to maximize differentiation between facets, without sacrificing coverage (i.e., some areas may 'fade out' with increased differentiation). When in doubt, aim for maximal coverage (facet differentiation/delineation will be addressed in step #8).

Note: this process is image-dependent; there are no set values. Threshold adjustment and visual confirmation of the facets gives the best results.

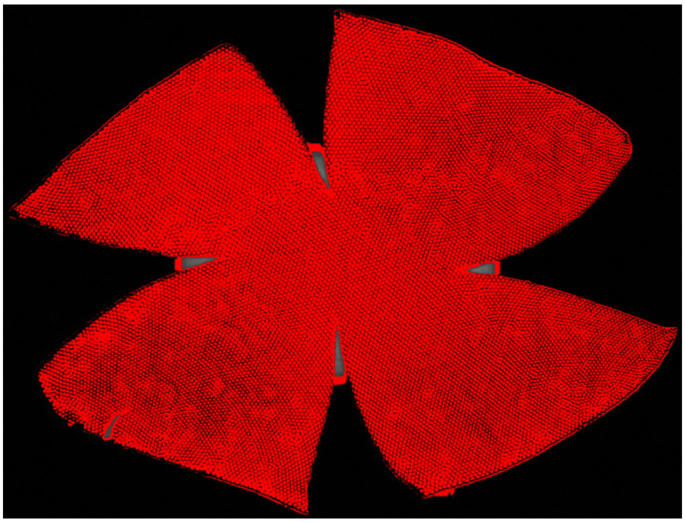

**8)** Click *Apply* when a satisfactory level of thresholding is achieved.

**9)** *Process* → *Binary* → *Watershed* (for individual facet differentiation/delineation).

a. The length of this process varies for each image/threshold level.

**10)** *Analyze* → *Analyze Particles*.

a. Size (mm^2): 0.0001-0.0009*
b. Circularity: 0.50-1.00
c. Show: Bare Outlines
d. Select Summarize

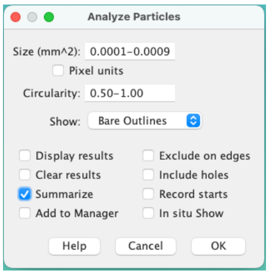


*Note: values were determined by measuring individual facet diameters to obtain a range of expected sizes. This may differ for every microscope and magnification level.

**11)** Click *OK*.

a. A summary table will appear, reporting the count.

b. Other variables may also be reported, but these are largely irrelevant for this process (e.g., corneal area will be measured differently).

c. A second window of the eye/cuticle will also appear, illustrating the bare outlines of the facets that were counted. This serves as a visual check of the process.

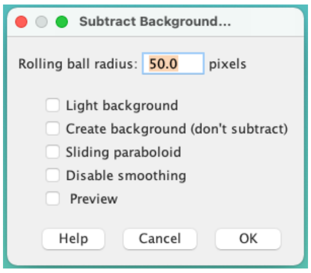

*Note: minor imperfections in the counts (bare outlines) may arise and are largely unavoidable. The goal is to minimize inconsistencies between samples.

**12)** Close the bare outlines and thresholded images, but DO NOT close the summary table.

**13)** Open the next image and repeat steps 4-12.

### Tibia length:

**1)** Open ImageJ/Fiji.

**2)** Open the desired image via *File* → *Open* or by dragging the image onto the console.

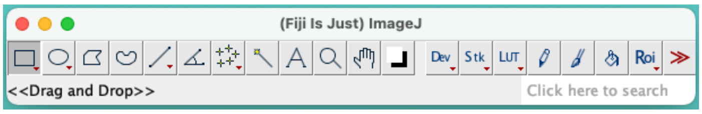


**3)** *Analyze* → *Set Scale* to specify the **distance in pixels** per **unit of length**. This is can be assessed via the line tool in the console.

Line tool

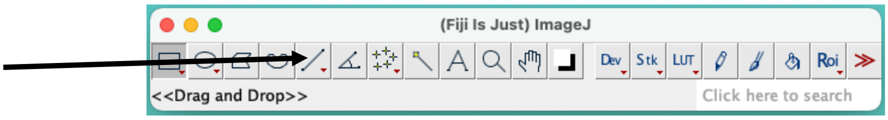


a. Click and drag a line the length of a known distance in the image, e.g., the scale bar, and then measure with *Analyze* → *Measure*.
b. Enter this value as the **Distance in pixels**, specify the **Known distance** (of the scale bar), and the **Unit of length**.
c. Select **Global** to apply this to all pictures opened in the current session.

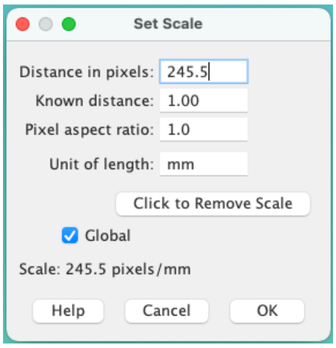


*Note: these values will need to be reassessed for images taken at different magnification levels.

**4)** Using the line tool as in step #3, draw a line along the length of the tibia.

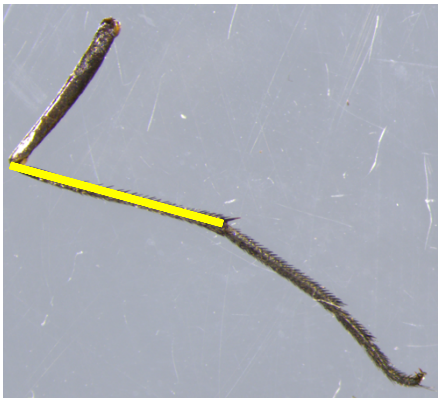

**5)** *Analyze* → *Measure*.

   a. Output variables in the summary table can be modified via *Analyze* → *Set Measurements*.

**6)** Close the image (<u>DO NOT</u> close the summary table), open a new image, and repeat steps 4-5.
